# High catalytic rate of the cold-active *Vibrio* alkaline phosphatase depends on a hydrogen bonding network involving a large interface loop

**DOI:** 10.1101/2020.10.27.357921

**Authors:** Jens Guðmundur Hjörleifsson, Ronny Helland, Manuela Magnúsdóttir, Bjarni Ásgeirsson

**Author notes:** Corresponding author: (JGH). **Structure data sets deposits:** PDB: 6QSQ.

## Abstract

The role of surface loops in mediating communication through residue networks is still a relatively poorly understood part of cold-adaptation of enzymes, especially in terms of their quaternary interactions. Alkaline phosphatase (AP) from the psychrophilic marine bacterium *Vibrio splendidus* (VAP) is characterized by an analogous large surface loop in each monomer, referred to as the large-loop, that hovers over the active site of the other monomer. It presumably has a role in VAP high catalytic efficiency that accompanies extremely low thermal stability. We designed several different mutagenic variants of VAP with the aim of removing inter-subunit interactions at the dimer interface. Breaking the inter-subunit contacts from one residue in particular (Arg336) caused diminished temperature stability of the catalytically potent conformation and a drop in catalytic rate by a half. The relative B-factors of the R336L crystal structure, compared to the wild-type, confirmed increased surface flexibility in a loop on the opposite monomer, but not in the large-loop. Contrary to expectations, the observed reduction in stability with an expected increase in dynamic mobility resulted in reduced catalytic rate. This contradicts common theories explaining high catalytic rates of enzyme from cold-adapted organisms as being due to reduced internal cohesion bringing increased dynamic flexibility to catalytic groups. The large-loop increases the area of the interface between the subunits through its contacts and may facilitate an alternating structural cycle demanded by a half-of-sites reaction mechanism through stronger ties, as the dimer oscillates between high affinity (active) or low phosphoryl-group affinity (inactive).

## Introduction

Enzymes are often adapted to a relatively narrow temperature range that compliments the needs of the host and/or the ambient temperature in a manner that also affects structural stability [1–3]. In order to understand the interplay between thermal stability and how temperature affects function, information on the physical chemistry that contributes to these properties must be collected. In addition, this knowledge can also have practical benefits for improving enzymes for use in industry [4, 5]. Previous studies have indicated that thermal adaptation depends on the particular protein under study, determining within protein families which structural features and bonding interactions are specifically of critical importance as determinants of stability or function [6, 7]. Due to the large number of non-covalent interactions within protein structures, in addition to the complex interactions with the solvent, the book-keeping of total energetics in the functional cycle of a protein becomes difficult [8]. This is not least due to the fact that the free energy between the folded and unfolded states is generally small compared with the total energy budget, when considering effects on stability. Furthermore, relating temperature to function, changing the temperature can alter which step becomes the rate-limiting step in the reaction pathway by specific effects on cohesive forces within the protein structure. Thus, one must be vigilant as to which step in the catalytic cycle is represented by *k*_cat_ when studying enzyme kinetics as a function of temperature. Changing the thermal energy in such a system can influence either the chemical events, conformational movements, or both [9].

The general plasticity of proteins is a key element of enzyme catalysis and the motions necessary for catalysis are an intrinsic property of each enzyme [10, 11]. The characteristic enzyme motions detected during catalysis are already present in the free enzyme with frequencies corresponding to the catalytic turnover rates. The turnover rate may be primarily determined by such motions rather than being limited by rates of the chemical steps [12]. This is particulary true for efficient enzymes working close to diffusion limits. Cold-adapted enzymes are ideal to test this paradigm, since they are usually highly homologous to their mesophilic counterparts, having the same basic structure and set of catalytically active site residues. Yet, their catalytic rate can be an order of magnitude higher as compared with more heat-tolerant variants. One debate relating to cold-adapted enzymes revolves around the question if they generally have increased global flexibility rather than local flexibility and, thus, have lower global thermal stability as a result of fewer non-covalent interactions internally. Given that increased flexibility is only needed in certain loops or domains that are directly related to catalysis, rigidity may be kept unchanged or even increased in other domains for folding stability [4, 6, 11–17]. Still, for some enzymes, a higher catalytic rate has been attributed to higher global flexibility at lower temperatures [18, 19]. Furthermore, the effect of allosteric effectors on enzyme dynamics is in many cases global [20]. Most of the literature has come to the conclusion that local flexibility is needed in certain domains, or around the active site, while other less catalytically relevant domains can be kept more rigid [21–24]. Thus, the local flexibility can be uncoupled from the global thermal stability and correlated to activity [25–27]. Comparison of B-factors in crystal structures between psychrophilic and mesophilic structures, as well as molecular dynamics (MD) analyses, have indicated participation from local flexibilities [17, 26, 28–35]. However, in some cases no differences in flexibility have been observed when comparing a psychrophilic and a mesophilic counterpart, such as for trypsin [36, 37] and citrate synthase [38]. Recently, a 2 μs MD simulation of endonuclease A showed no difference in flexibility between the cold-adapted strain from *Aliibrio salmonicida* and the mesophilic strain *Vibrio cholera* [39].

The fact that flexible local areas of functional importance can in some cases be decoupled from the global stability of the enzyme holds promise of explaining the use of the often higher entropic state of loops to improve catalytic efficiency in cold-adapted enzymes [32, 40, 41]. Here, we turned our attention to a long and unstructured interface region of an alkaline phosphatase (AP) previously isolated and characterized from a *Vibrio splendidus* bacterium (VAP, UniProtKB Q93P54) [42]. VAP, together with *Halomonas* AP and *Cobetia marina* AP, has the highest reported turnover rate for APs in the literature and all three contain the same large interface loop (referred to in this paper as the large-loop) [43, 44]. In VAP, the loop structure has twelve intra-molecular hydrogen bonds within the loop where six of those are between main chain atoms. The loop interface towards the other subunit consists of eleven inter-molecular bonds where most are located near the stem where the loop starts and ends. The main hydrogen bonding region connecting the loop to the other subunit is located around Arg336. In this work, we mutated this arginine to leucine and also made variants where its binding partners were changed to rupture interacting hydrogen bonds. Results indicated that the loss of non-covalent bonds at the loop interface resulted in increase in flexibility (higher relative B-factors) which contributed to a large decrease in stability in some cases, and interestingly in decreased catalytic activity where the opposite might be expected.

## Methods

### Materials

Chemicals were obtained from Sigma-Aldrich (Schnelldorf, Germany) or Merck (Darmstadt, Germany) with the following exceptions. Strep-Tactin Sepharose, (2-(4’-hydroxy-benzene-azo) benzoic acid (HABA), desthiobiotin and anhydrotetracycline (AHTC) were from IBA GmbH (Germany). Bacto Yeast extract was obtained from Becton, Dickinson and Company (France). Primers were obtained from TAG (Copenhagen, Denmark). Pfu polymerase, DnpI nuclease and PageRuler protein ladders were from ThermoFisher Scientific.

### Generation of VAP variants and protein production

Mutagenesis was performed on the VAP gene as described elsewhere using the QuikChange method [45]. Protein expression and purification protocols have also been described in detail previously [45].

### VAP enzyme activity assay

Standard assays during purification were performed with 5.0 mM *p*-nitrophenyl phosphate (*p*-NPP, biscyclohexylamine salt) in 1.0 M diethanolamine with 1.0 mM MgCl_2_ at pH 9.8 and 25 °C. The absorbance of the reaction product was measured at 405 nm over a 30-s period in a temperature regulated Thermo spectrophotometer. An extinction coefficient of 18.5 M^−1^cm^−1^ was used at pH 9.8 for the calculation of international enzyme units and conversion into *k*_cat_ values performed using a calculated extinction coefficient at 280 nm for VAP of 61310 M^−1^ cm^−1^.

### Determination of kinetic constants

Steady-state kinetics were performed at 10°C in 0.1 M Caps, 1.0 mM MgCl_2_, 0.5 M NaCl at pH 9.8. The pH was corrected for temperature variation. The substrate (pNPP) was diluted in the range 0.0 mM to 1.0 mM. Rate constants were obtained by direct fit to the Michaelis-Menten equation using the software GraphPad Prism 6^®^ (San Diego, California, USA) or Kaleidagraph® (Reading, Pennsylvania, USA).

### Thermal stability measurements

Determination of *T*_50%_, defined here as the temperature needed to reduce initial activity by 50% over a 30 min period, was performed by incubating an enzyme sample in a glass tube containing pre-heated buffer in the range of 10-27°C without NaCl and 50-60°C with 0.5 M NaCl present (which stabilizes the very labile local structural in the active site) [46]. Experiments were initiated by adding the enzyme (10-15 μl) to the medium (500-1000 μl). The rate constant at each temperature was determined by regularly withdrawing aliquots and following the reduction in activity using the standard activity assay at 25°C. From an Arrhenius plot (ln*k* vs. 1/*T*), the activation energy (*E*_a_) was obtained. *T*_50%_ was calculated from the Arrhenius equation, where the *k* (s^−1^) for 50% loss of activity after 30 min was used in each case (*k*=ln 2/30 min · 60 s/min^−1^) to give *T*_50%_ = (*E*_a_ × 1000)/*R*(ln*A*-ln*k*). Samples for *T*_m_ determination, as a measure of secondary structure stability, were dialyzed overnight in 25 mM Mops, 1 mM MgSO_4_ at pH 8.0 and absorbance at 280 nm measured to determine protein concentration. A JASCO J-810 circular dichroism (CD) spectropolarimeter (Tokyo, Japan) was used to determine melting-curves at 222 nm over the temperature range 20-90°C in a 2 mm cuvette with a rise in temperature of 1°C/min. For monomer unfolding, a two-state pathway was assumed (*N*⇄*D*, where *N* is native the state and *D* is the denatured state). The original traces were normalized, and the *T*_m_ determined at *F*_U_ = 0.5 by a direct fit to a sigmoid curve using the software Kaleidagraph®.

### X-ray crystallography

Crystallization conditions were screened for using a Phoenix crystallization robot (Art Robbins Instruments) by the sitting-drop vapor-diffusion method. About 400 commercial and in-house made conditions were screened by using MRC plates with a 60 μl reservoir solution per well, and drop solutions were prepared by mixing 0.25 μl well solution and 0.25 μl protein solution at 12 mg/ml (20 mM Tris, 1 mM MgSO_4_, 5 mM MgCl_2_, 1.3 mM desthiobiotin, 0.25 M NaCl, 15% (v/v) ethylene glycol, pH 8.0). Crystals were obtained from 20% PEG 3350 (w/v) and 0.2 M potassium formate.

Data to 2.0 Å was collected at the BL14.1 synchrotron radiation station of Bessy (Berlin, Germany). Data was integrated and scaled using XDS [47] and AIMLESS [48, 49]. The structure of VAP R336L was solved by molecular replacement using MOLREP [50] of the CCP4 suite [49] and native VAP (PDB ID: 3E2D) [28] as template. Inspection of electron density maps were carried out in Coot [51], followed by positional refinement in REFMAC [52]. Data collection and refinement statistics are listed in Table 2.

## Results

### The structure of the large-loop of VAP

Fig 1 shows the crystal structure of the wild-type VAP (*Vibrio* alkaline phosphatase) focusing on the intra-molecular interactions to residues surrounding R336 on the large interface loop (in red) and the inter-subunit interaction connecting the loop to the other subunit, detailed below. Several structure alignments of VAP with other AP crystal structures highlight the differences in lengths of the interface loop. In fact, the *E. coli* variant (ECAP) does not harbor the VAP large-loop, but has an N-terminal interface loop followed by a short helix instead. Same goes for the mammalian phosphatases which all have an N-terminal interface helix, as does the shrimp AP. On the other hand, the Antarctic AP (from the bacterium strain TAB5) is shown to have a similar large-loop insert as VAP. However, in this case, it resembles more closely a β-structure and is directed away from the interface and active site in the crystal structure (likely a crystal-packing artifact) [53]. Another highly homologous structure to VAP is the *Shewenella* AP with a much shorter interface loop. Finally, the AP from the halophilic bacterium *Halomonas* has near-identical structure to VAP, including the large-loop.

**Fig 1.**
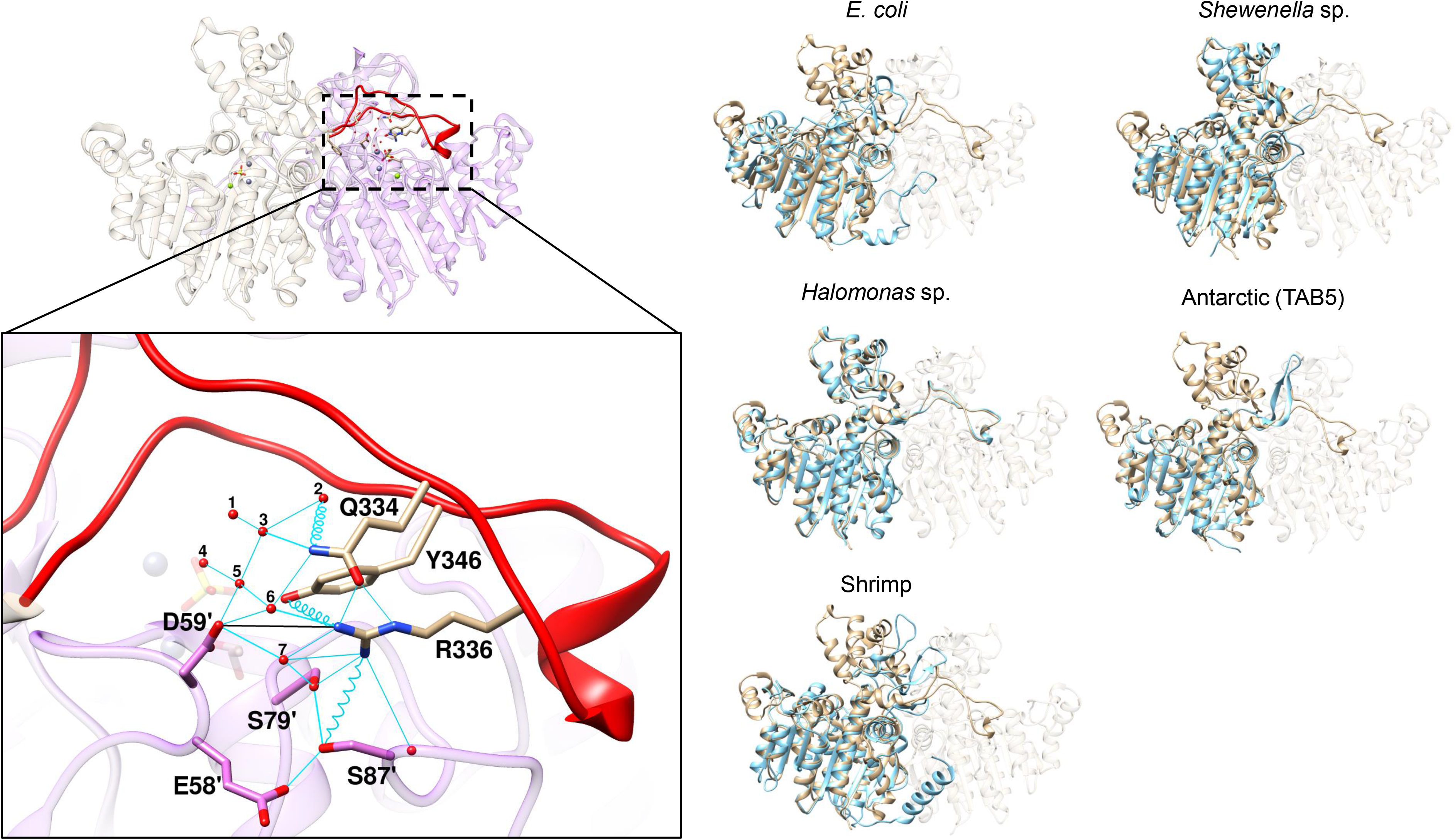
The large-loop in *Vibrio* alkaline phosphatase (VAP). Left panel: The inset shows the inter- and intra-molecular interactions surrounding R336 in the large interface loop of VAP (PDB ID: 3E2D). Key water molecules are numbered. Hydrogen bonds that are < 3.0 Å are shown as cyan lines and weak hydrogen bonds (3.0-5.0 Å) are shown as cyan springs. Salt-bridges are denoted by a black line and Zn^2+^ and Mg^2+^ ions are represented as navy-blue and bright-green, respectively (seen transparent in the background to emphasize the location of the active site). Residues from monomer B are denoted by a mark symbol (‘). Right panel: VAP (in tan color) was aligned to the following AP structures (in cyan) to compare large-loop regions: *E. coli* (PDB ID: 3TG0), *Shewenella* sp. (PDB ID: 3A52), *Halomonas* sp. (PDB ID: 3WBH), Atlantic shrimp (PDB ID: 1SHN) and Antarctic bacterium TAB5 (PDB ID: 2W5W).

### Replacement of the large-loop residue Arg336 affects both activity and stability of VAP

For VAP, an Arg residue at position 336 is a central residue in forming and maintaining extensive intra-chain (with Q334 and Y346) and inter-subunit (with S79 and S87 at subunit B) hydrogen-bonding network (Fig. 1, inset). Additionally, attached to these residues is an extensive network of structured water molecules which form a hydrogen-bond network. Furthermore, a long-range inter-chain salt-bridge (4.9 Å) exists between D59 of subunit B and R336 of subunit A, and between D59 of subunit A and R336 of subunit B. To determine the importance of these interactions, we produced the R336L variant and tested its effect on protein stability and activity (Table 1). The Arg to Leu mutation was selected to prevent any side-chain mediated polar interactions, and at the same time cause only a minimal change in the side-chain’s steric interactions. Additional variants were also produced to see the effect of spoiling individual hydrogen bonds. Q334 and Y346 are on the large-loop on either side of R336, whereas Ser79 and Ser87 are part of the opposite subunit (see below). We have previously shown that during heat inactivation the wild-type enzyme is transformed to an inactive dimer intermediate [54]. The stability of the weakest part of the VAP structure that the activity depends on, and is particularly sensitive to temperature, can be quantified as the rate of inactivation, encapsulated in the parameter *T_50%_* (the temperature that will give the 50% inactivation during a 30 min incubation at that particular temperature). The global denaturation of the secondary structures was monitored by CD spectroscopy to give T_m_.

**Table 1.**
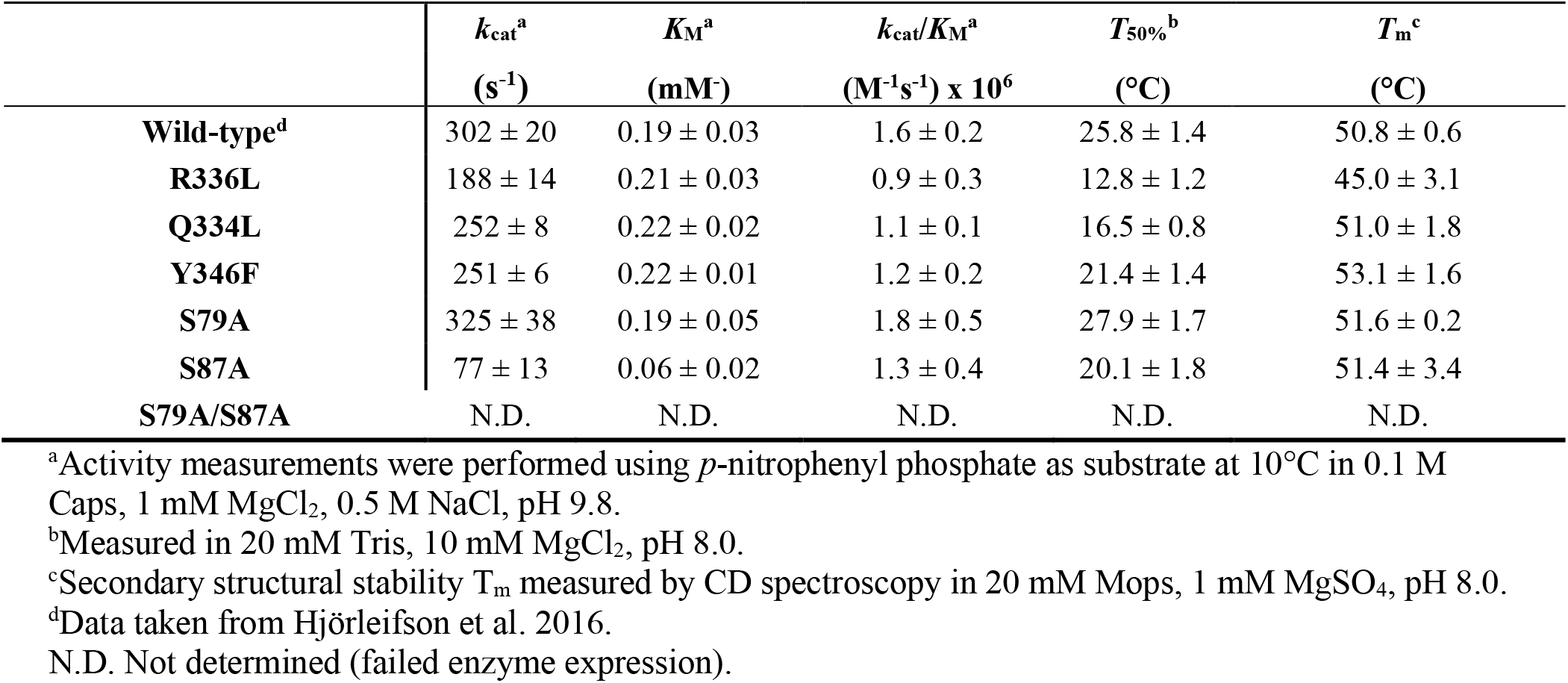
Temperature stability and kinetic constants of VAP and relevant variants. The results are presented as mean ± SD from repeated experiments.

| | $k_{cat}^a$<br>(s<sup>-1</sup>) | $K_M^a$<br>(mM) | $k_{cat}/K_M^a$<br>(M<sup>-1</sup>s<sup>-1</sup>) x 10<sup>6</sup> | $T_{50\%}^b$<br>(°C) | $T_m^c$<br>(°C) |
| --- | --- | --- | --- | --- | --- |
| **Wild-type<sup>d</sup>** | 302 ± 20 | 0.19 ± 0.03 | 1.6 ± 0.2 | 25.8 ± 1.4 | 50.8 ± 0.6 |
| **R336L** | 188 ± 14 | 0.21 ± 0.03 | 0.9 ± 0.3 | 12.8 ± 1.2 | 45.0 ± 3.1 |
| **Q334L** | 252 ± 8 | 0.22 ± 0.02 | 1.1 ± 0.1 | 16.5 ± 0.8 | 51.0 ± 1.8 |
| **Y346F** | 251 ± 6 | 0.22 ± 0.01 | 1.2 ± 0.2 | 21.4 ± 1.4 | 53.1 ± 1.6 |
| **S79A** | 325 ± 38 | 0.19 ± 0.05 | 1.8 ± 0.5 | 27.9 ± 1.7 | 51.6 ± 0.2 |
| **S87A** | 77 ± 13 | 0.06 ± 0.02 | 1.3 ± 0.4 | 20.1 ± 1.8 | 51.4 ± 3.4 |
| **S79A/S87A** | N.D. | N.D. | N.D. | N.D. | N.D. |

<sup>a</sup>Activity measurements were performed using *p*-nitrophenyl phosphate as substrate at 10°C in 0.1 M Caps, 1 mM MgCl<sub>2</sub>, 0.5 M NaCl, pH 9.8.
<sup>b</sup>Measured in 20 mM Tris, 10 mM MgCl<sub>2</sub>, pH 8.0.
<sup>c</sup>Secondary structural stability $T_m$ measured by CD spectroscopy in 20 mM Mops, 1 mM MgSO<sub>4</sub>, pH 8.0.
<sup>d</sup>Data taken from Hjörleifson et al. 2016.
N.D. Not determined (failed enzyme expression).

The R336L variant displayed a significant decrease in *k*_cat_ (62% of wild-type), whereas the *K*_M_ was practically unchanged (Table 1). Concurrently, the *T*_50%_ was greatly decreased from 25.8°C for the wild-type to 12.8°C in R336L. When the buffer medium contained an addition of 0.5 M NaCl, the R336L variant had the same *T*_50%_ as the wild-type, close to 53 °C as previously reported [46]. The *T_m_* was slightly decreased for R336L compared to wild-type, in stark contrast to the drastic difference in temperature tolerance of the active form of the enzyme.

### Loop flexibility as reflected by B-factors in the R336L structure

The crystal structure of R336L VAP variant was solved at 2.0 Å resolution (PDB ID: 6QSQ, statistics in Table 2). The wild-type and R336L VAP structures were overall similar, although the crystal packing was different (P 2_1_ 2_1_ 2 and C 2 2 2_1_ space group, respectively). Some side chains in relatively open parts of the structure displayed slight changes in conformation. The asymmetric unit for the R336L structure was the monomer, and this was translated into the biological unit for R336L as a dimeric structure. One sulphate ion and one desthiobiotin molecule (picked up from the elution off the StrepTactin affinity column) were identified in the electron density, in addition to the catalytic Zn and Mg ions. The asymmetric unit for the previously solved crystal structure of the wild-type VAP was the dimer [28], and a PEG molecule was observed in the same position as the desthiobiotin. The only significant structural difference between the wild-type and the R336L variant was the rotation of the side chain at position 88, where E88 of chain B is oriented away from R336 of chain A in the wild-type, whereas it is pointed towards the replacement residue in the variant L336.

**Table 2.**
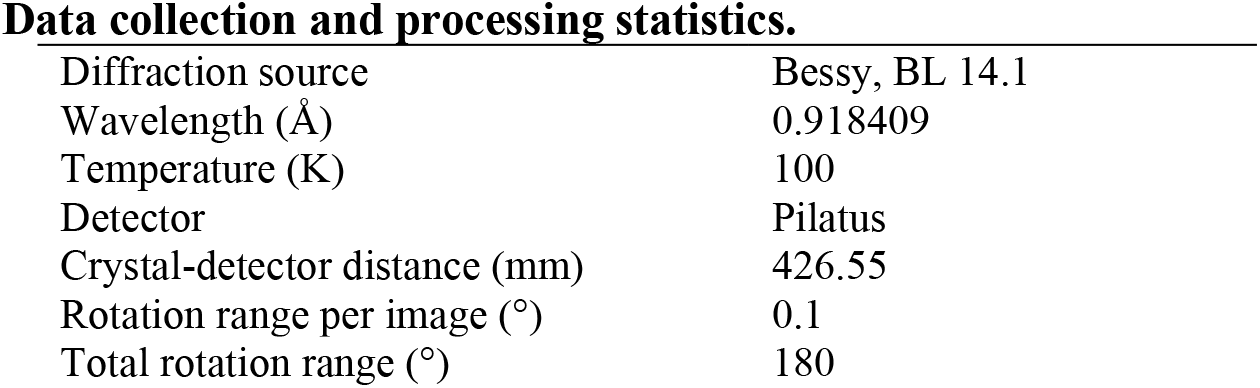
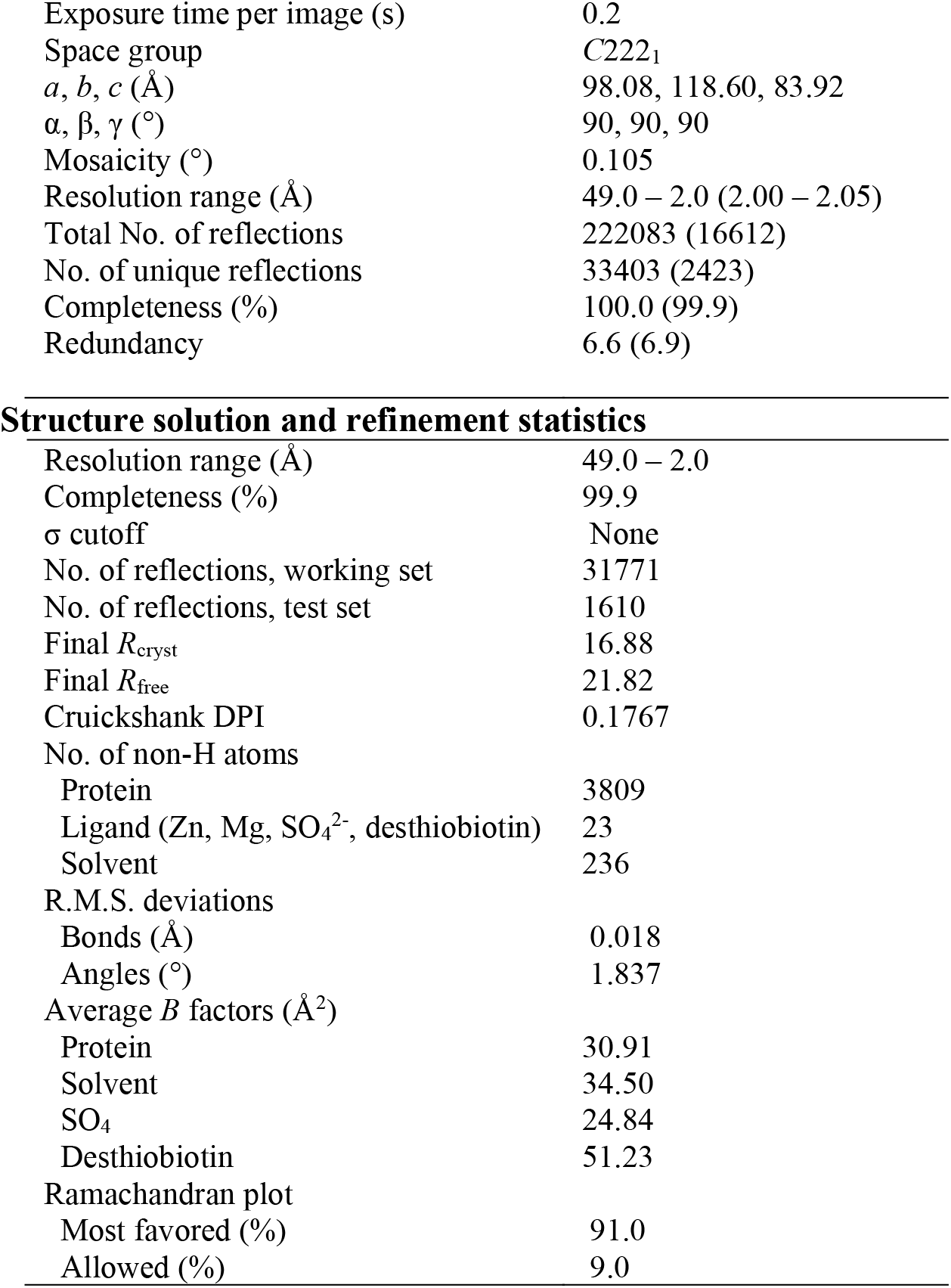
X-ray crystallographic data for the R336L VAP structure (PDB: 6QSQ). Values for the outer shell are given in parentheses.

Data collection and processing statistics.
|  |  |
| --- | --- |
| Diffraction source | Bessy, BL 14.1 |
| Wavelength (Å) | 0.918409 |
| Temperature (K) | 100 |
| Detector | Pilatus |
| Crystal-detector distance (mm) | 426.55 |
| Rotation range per image (°) | 0.1 |
| Total rotation range (°) | 180 |
| Exposure time per image (s) | 0.2 |
| Space group | C222<sub>1</sub> |
| *a*, *b*, *c* (Å) | 98.08, 118.60, 83.92 |
| $\alpha$, $\beta$, $\gamma$ (°) | 90, 90, 90 |
| Mosaicity (°) | 0.105 |
| Resolution range (Å) | 49.0 – 2.0 (2.00 – 2.05) |
| Total No. of reflections | 222083 (16612) |
| No. of unique reflections | 33403 (2423) |
| Completeness (%) | 100.0 (99.9) |
| Redundancy | 6.6 (6.9) |
| Structure solution and refinement statistics |  |
| Resolution range (Å) | 49.0 – 2.0 |
| Completeness (%) | 99.9 |
| $\sigma$ cutoff | None |
| No. of reflections, working set | 31771 |
| No. of reflections, test set | 1610 |
| Final $R_{\text{cryst}}$ | 16.88 |
| Final $R_{\text{free}}$ | 21.82 |
| Cruickshank DPI | 0.1767 |
| No. of non-H atoms |  |
| Protein | 3809 |
| Ligand (Zn, Mg, SO<sub>4</sub><sup>2-</sup>, desthiobiotin) | 23 |
| Solvent | 236 |
| R.M.S. deviations |  |
| Bonds (Å) | 0.018 |
| Angles (°) | 1.837 |
| Average *B* factors (Å<sup>2</sup>) |  |
| Protein | 30.91 |
| Solvent | 34.50 |
| SO<sub>4</sub> | 24.84 |
| Desthiobiotin | 51.23 |
| Ramachandran plot |  |
| Most favored (%) | 91.0 |
| Allowed (%) | 9.0 |

The average main-chain root mean square deviation (RMSD) between the wild-type monomer A and the R336L variant was 0.35 Å indicating a near-perfect rotational symmetry. However, a correlation was observed in the traces for main-chain asymmetry within the dimeric structures in certain regions (Fig 2A). This was evaluated by superimposing each monomer A on monomer B following distance measurements of respective C_α_ atoms. Since the asymmetric unit was the monomer for the R336L structure, such analysis could not be performed. Instead the R336L monomer was superimposed onto wild-type monomers A and B, respectively. The largest difference around residue 225 was due to asymmetric binding of a sulphate ion to monomer A in the wild-type crystal structure, which is not present either in the wild-type monomer B or the R336L structure. The loop region of residues 153-183, 325-353 and 365-395 can all be considered genetic inserts, when compared to the *E. coli* AP counterpart, and we have previously designated those as inserts I, II and III [28]. These insert regions all showed relatively higher RMSD values than other regions, albeit with RMSD of < 1.0 Å. This is in an accordance with an MD study, previously performed on VAP, where the insert regions were shown to have high main-chain asymmetry due to dynamic mobility [35]. Interestingly, a highly homologous structure of AP from the salt-adapted extremophile *Halomonas* sp. showed a similar trend in the corresponding insert regions (denoted by arrows in Fig. 2). We believe that these observations are significant, and not due to crystal contacts in the unit cells, since the three evaluated structures (PDB ID: 3E2D, 6QSQ and 3WBH) were all solved from crystals with widely different space groups (thus, not sharing the same crystal contacts). However, we do like to note that the higher main-chain RSMD values in these regions might arise from poorer quality of electron density in high mobility regions.

**Fig 2.**
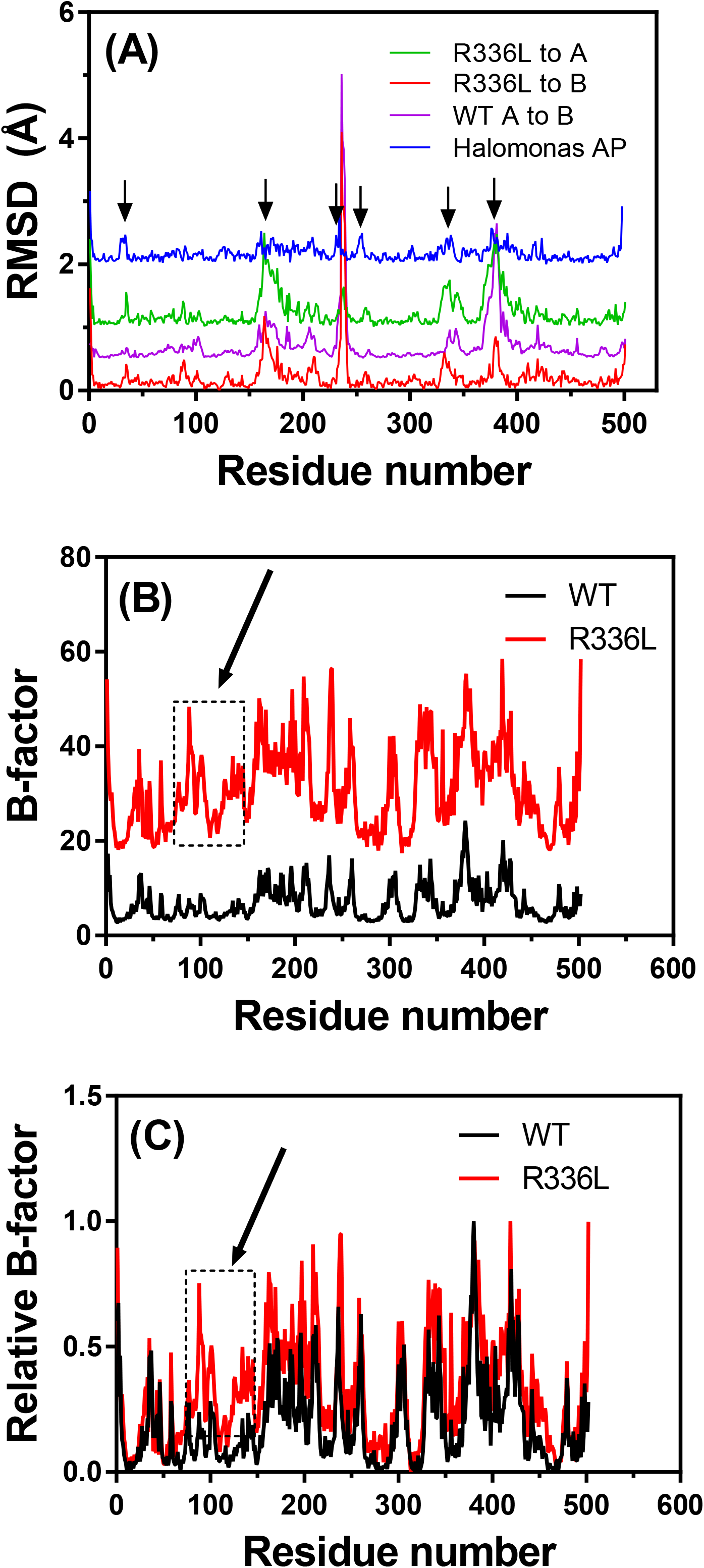
Comparison of main-chain dynamics in the wild-type and R336L VAP crystal structures. (A) Main-chain root-mean-square-deviation (RMSD) of the wild-type monomer A superimposed onto monomer B (in purple), the R336L structure superimposed onto wild-type monomer A (in green) and wild-type monomer B (in red). The average RMSD was 0.30 Å and 0.35 Å for wild-type and R336L, respectively. As for comparison, monomers A and B of the *Halomonas* sp. AP main-chain RMSD are shown (in blue). Arrows denote areas with correlated differences in the main-chain positions indicating main-chain asymmetry during X-ray data acquisition. Note that the data traces have been nudged for clarity. (B) Raw temperature B-factors of the wild-type (black) and R336L (red) VAP structures. (C) The B-factors from (B) normalized to the highest B-factor value in each data set. An increase in mobility (B-factor) can be observed in the area labelled by an arrow (residues 70-150).

The absolute B-factor values for the wild-type and R336L structures can be seen in Fig. 2B. The three insert regions associated with extra loops in VAP that were noted above, all give spikes in B-factor indicating disorder within the crystal. However, by comparing main chain RMSD values to B-factors in Figs. 2A and 2B, respectively, no clear correlation between spikes in RMSD values and B-factors could be established. By normalizing the B-factors for the wild-type and R336L structures, an overlay comparison fitted well in most regions except in the residue regions 81-100 and 115-150 (denoted by an arrow in Figs. 2B and 2C). A “worm” rendering of the normalized B-factors in the large-loop region (Fig. 3) showed that on average the structure is most disordered in the vicinity of the residue replacement R336L (Figs. 3A and 3B). It should be noted that the region containing residues 65-150 forms the helix on which the S65, the nucleophile in the active site, is located. Residues S79 and S87 are inter-hydrogen bonded to R336 (from the other monomer) and the helix-turn-helix motif where R129 is positioned. R129 has a very important role as the main phosphate-binding residue for incoming substrates as well as the phosphate ion product. Thus, increased dynamics in this region might be expected to affect the catalytic rate through facilitated release of the product (this being the rate-limiting step at alkaline pH). Interestingly, higher relative B-factors were not observed for the main chain of the large-loop itself (residues 324-354, Fig. 2C), which indicates that the loop remains well anchored by other interactions when the R336 connections are broken, and the primary result of the R336L mutation is the long-range propagation of a dynamical effect. The distance of this information transfer can be considered to span 60 Å since that is the distance between the active sites of each monomer. It may be noted, that despite more dynamic mobility in the active site that was observed in the R336L variant (by T_50%_), the measured K_M_ value for p-nitrophenyl phosphate remained practically unchanged. The interaction with substrate in alkaline phosphatases is dominated by binding between the phosphoryl group to the metal ions and Arg129, and the mutational changes do not seem to affect these.

**Fig 3.**
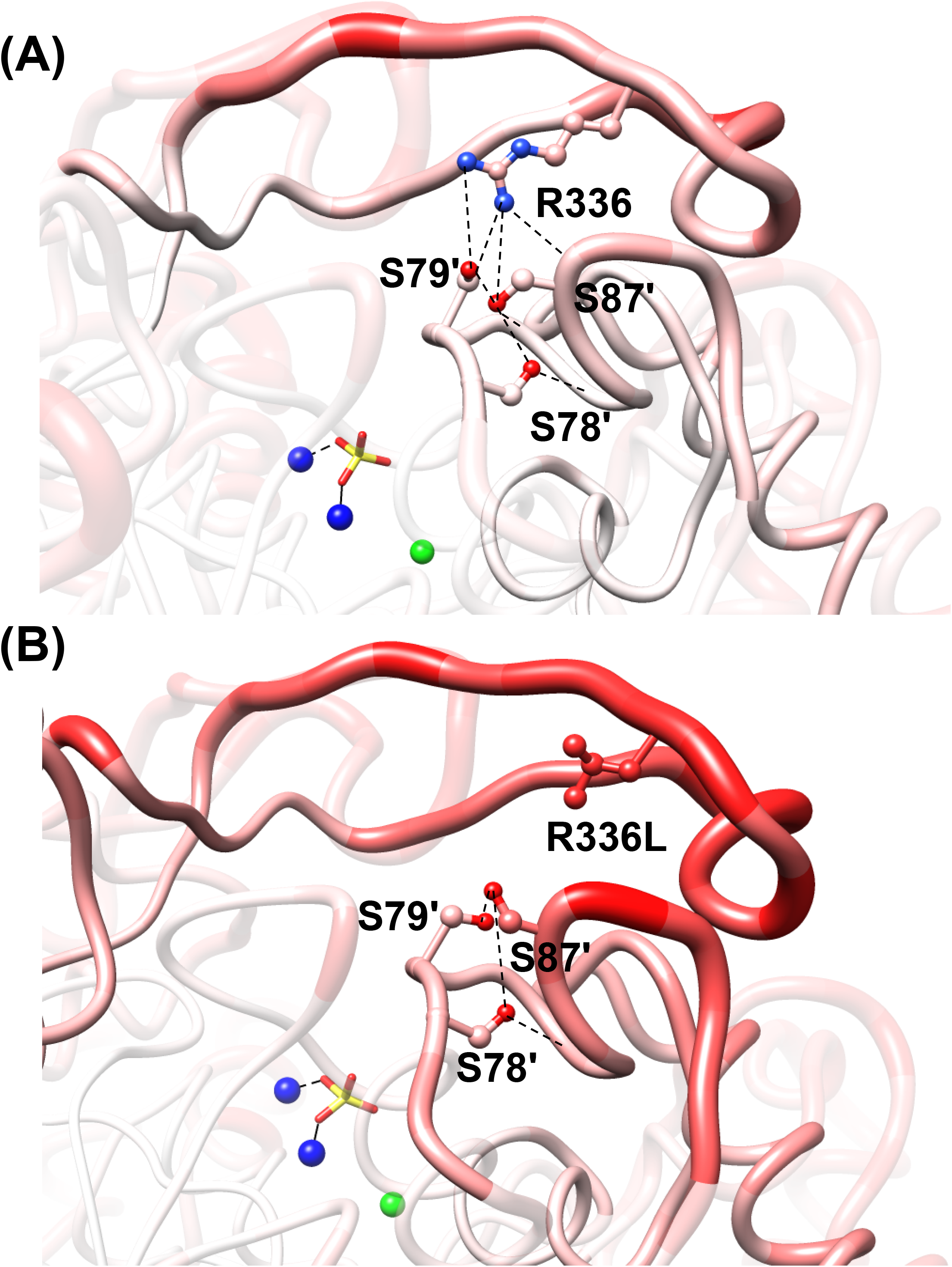
A main-chain B-factors analysis of the large-loop region of wild-type VAP (A) and the variant R336L (B). The thickness of the “worms” goes from 0.3 Å to 1.25 Å for the lowest 5% to the highest 5% of the B-factors, respectively. Additionally, the main-chain and side-chains are colored from blue (0.3 Å) to red (1.25 Å) with white as a mid-score (0.80 Å). Hydrogen bonds are shown as broken black lines, and Zn^2+^ and Mg^2+^ as blue and green spheres, respectively.

### The role of large-loop intra-chain and inter-chain hydrogen bonds

The R336L residue replacement leads to a loss of many intra-molecular hydrogen bonds within the loop structure and also ruptures inter-subunit hydrogen bonds with S79 and S87. Several variants were produced to locate various effects more precisely and quantify the effect of reducing these intra-molecular and inter-molecular hydrogen bonds. The Y346F and Q344L variants should reduce intra-molecular hydrogen bonds with R336 by one and two, respectively, as well as generate a loss of an additional hydrogen bond with structured water (Fig. 1). Table 1 shows, that the Y346F and Q334L variants had similar catalytic properties as the wild-type (k_cat_ and K_M_) but both showed a decrease in *T*_50%_ from 25.8°C (wild-type) to 21.4°C and 16.5°C, respectively. We relate this to the most vulnerable part of the enzyme where interactions are loosest and dynamic mobility greatest, namely the active site. In contrast, the *T*_m_ was unchanged, and this we relate to the global structural stability involving all weak interactions in the molecule and bonding between the two monomers. The *T*_50%_ was less affected by the reduction of inter-subunit hydrogen bonds (R336L, S79A, S87A) than by reducing only intra-molecular hydrogen bonds within the loop structure (Q334L and Y346F). In all cases, except for S79A, there was a significant decrease in *T*_50%_. By producing the double variant S79A/S87A, we would alter the inter-subunit hydrogen-binding partners of R336L in the opposite subunit. However, we were unable to produce the S79A/S87A in the folded form despite repeated effort.

## Discussion

We have shown experimentally that a reduction in the number of hydrogen bonds formed by R336 in the VAP large-loop caused a decrease in thermal stability (Table 1) and also resulted in lower *k*_cat_ with respect to the wild-type (Table 1). This suggests that removing the charge and hydrogen bonding potential of R336 can alter the conformational landscape of VAP's active residues, affecting conformational states that are important for catalysis. It has previously been shown, that the isozyme specific properties of mammalian APs canbe modulated by modifications in a flexible surface loop [55], or by affecting the local network of interactions close to the catalytic site [56]. More generally, surface loop flexibility has been shown to play a major role in the temperature adaptation of enzymes, by providing a way to replace enthalpic requirements in the total energy budget with entropy contribution stemming from more mobility, in particularly in surface loop [57]. Interestingly, in some cases where loop structures from thermophilic enzymes have been introduced to mesophilic counterparts, the stability was greatly increased, indicating topological roles of loops in stabilizing protein structures towards higher temperature [58].

All the APs that have this particular large-loop under study here, and have been kinetically characterized, are significantly more active than variants with it absent (or having a shorter form of it). Thus, we hypothesized that the large-loop has a role in increasing product release, as the rate-limiting step at neutral to alkaline pH in APs is release of inorganic phosphate [59–61]. Furthermore, we have shown that addition of 20% w/v sucrose decreased the activity VAP up to 50% indicating a diffusion dependent step, such as product release, as the rate-limiting step (Hjörleifsson and Ásgeirsson, 2017 supporting material) [46]. A lower k_cat_ from this cause might also be linked with a lower K_M_ value due to tighter binding of product and substrate (the phosphate moiety is the major determinant for binding affinity of both). Lower K_M_ value was only associated with the S79A variant, but K_M_ was unchanged for R336L. Thus, the k_cat_ decrease in R366L could only partially be coupled with higher substrate/product affinities. The new R336L crystal structure showed increased dynamics in loops (in terms of B-factors) close to the active site of the opposite subunit, albeit with no significant changes in main chain dynamics for the large-loop as a whole. A previous MD study pointed out a long-range communication link through a hydrogen bonding network from the large-loop (R336) to the catalytic cleft involving the D59 and E81 side-chains [35]. By disrupting the hydrogen-bonding cluster(s) mediated by R336 by replacement with leucine, we showed conclusively the importance of the long-range communication mediated by R336. However, it is at present difficult to conclude in finer detail how these changes in dynamics lead to a decrease in activity, although such dynamic couplings to catalytic cores have been shown to be mediated by surface loops [62]. Possibly, a build-up of interactions, involving hydrogen bond network(s) at the large-loop, is needed for a reciprocal conformational change of the enzyme subunits leading to the release of the product in each catalytic cycle. Reducing the number of participants in these hydrogen bonding interactions might slow down the conversions between the E·P (non-covalent bonded phosphate ion in active site) and E+P (non-covalent bonded phosphate with solvent components outside the active site) states due to less rigid coupling between the cooperating subunits. Increased flexibility of the loops would only be a partial contributor to motions that promoted faster release of product.

The R336L variant showed both a decrease in global stability in terms of *T*_m_ and in *T*_50%_, where the difference in *T*_50%_ was very large, from 25.8 ± 1.4 °C to 12.8 ± 1.2 °C, for wild-type and R336L respectively. This is consistent with a key role for the large-loop in providing thermal stability for the active conformation. We tested the effect of adding chloride ions, since they have been shown to facilitate efficient refolding of VAP when the enzyme was denatured in urea [54]. However, with 0.5 M NaCl present, there was no difference in *T*_50%_ for the wild-type nor the R336L variant. In fact, evidence suggests that halophilic- and psychrophilic adaptation has co-evolved in enzymes [63].

Individual inter-subunit hydrogen bonds with R336 were broken by producing the S79A and S87A variants. The S79A/S87A double variant was not successfully expressed in our *E. coli* expression system. Interestingly, the k_cat_ was not altered in the S79A single variant while there was a large decrease in the S87A variant. S87A was the only variant tested in the present study where a large decrease in K_M_ occurred. As discussed above, observing lower k_cat_ with lower K_M_ fits well with a rate-limiting product-release mechanism, being the result of tighter substrate and product binding.

The terms “flexibility” and “dynamics” have an uncertain meaning when it comes to using them to describe how an enzyme functions. Therefore, it has proven difficult to generalize when it comes to explaining the role flexibility in enzyme catalysis. Flexibility generally refers to the range of movement and how easily something can fluctuate through the possible intermediary states, while dynamics refer to how fast changes or fluctuations progress with time. The importance of local dynamics in cold-adaption has been reported for adenylate kinase where a clear role for dynamics in catalytic rate regulation was observed [64]. The dynamics in an AMP binding domain were correlated to changes in the turnover speed of the rate-limiting conformational change in the enzyme, mainly due to changes in entropy. Another important consideration that commonly gets left out of energy discussions related to enzyme function is the effect of water solvation on protein structures and interactions with substrates, where entropy weights in prominently and is clearly both involved in catalysis as well as the binding interaction with small substrate molecules and ions. In fact, in our case, there is a lot of structured water bound at the large-loop dimer interface of VAP with an unknown functional role. Isaksen et al. [15], proposed that protein-solvent interfacial surfaces differ between cold-adapted and thermophilic enzymes due to point mutations that disrupt surface hydrogen bonding networks and bound water. This was observed using calculations of “high-precision” Arrhenius plots and thermodynamic activation parameters. It is experimentally challenging to prove these theories, but recently Kim et al. [65] used time-resolved crystallography and NMR to show that release of bound water network during binding of substrate facilitated an entropy driven conformational change that was involved both in the binding of substrate and the release product in a homodimeric fluoroacetate dehalogenase. These accumulated results indicate that both inter- and intra-molecular bonds in loop structures of homodimeric enzymes can be highly important for stability and efficient catalytic activity.

## Conclusions

Overall, the inter-molecular interface in the VAP dimer between the large-loop of one subunit and two serine loop-residues in the opposite subunit is important for the catalytic cycle. R336 is at the heart of this inter-molecular hydrogen-bonded network that communicates large-loop interactions to the active sites. R336L is a prominent ionic residue on the large-loop that forms most of these inter-subunit bonds and affects catalytic competence. Further conclusions will await experimental confirmation of a reciprocal asymmetry in the dimer, where each monomer presumably oscillates conditionally between two main conformations in an opposite manner to its partner.

## Abbreviations

AP: Alkaline phosphatase
VAP: *Vibrio* alkaline phosphatase
MD: Molecular dynamics
*p*-NPP: *para*-nitrophenyl phosphate
CD: Circular dichroism
ECAP: *E. coli* alkaline phosphatase
RMSD: Root mean square deviation

## Data accessibility

The crystal structure of VAP R336L variant has been deposited and released in the protein data bank (PDB) with the PDB ID: 6QSQ.

## Author contributions

JGH and BA conceived and supervised the study; JGH and BH designed experiments; JGH, BA, MM and RH and performed experiments; JGH, BA, MM, and RH analyzed data; RH collected X-ray structure data and performed analysis; JGH and BA wrote the original draft manuscript; JGH, BA, MM and RH made manuscript revisions.

## Acknowledgments

Gratitude is extended to the Icelandic Research Fund (grant number 060219021-3 and 141619-05-3) and the University of Iceland Research Fund for supporting this project financially. The “Syncnøyt programme” of the Research Council of Norway (project # 247732) is gratefully acknowledged for financial support, as is the provision of beamtime at BESSY beamline 14, at the electron storage ring operated by the Helmholtz-Zentrum, Berlin.

